# A mechanistic model of threat-driven respiration and its homeostatic recovery in anxiety and depression

**DOI:** 10.64898/2026.10.16.759543

**Authors:** Malik Faizan, P. J. White, Indrakshi Dey

## Abstract

Anxiety and depression change how people breathe, often early in the illness, and wearable sensors now allow breathing to be monitored continuously. This work presents, to our knowledge, the first mechanistic model of how the brain’s threat response sets breathing rate and depth and how fast breathing recovers after stress. A threat signal from the amygdala, shaped by stimulus magnitude and by cognitive appraisal (coping capacity and perceived stress), drives the breathing-control center that receives lung feedback: stretch receptors slow breathing and airway receptors speed it up. A simple electrical circuit stores the threat signal and expresses recovery as one time constant. We prove the model’s key properties, test them under noise and parameter variation, and compare it with chest-belt recordings from 15 adults in the Wearable Stress and Affect Detection (WESAD) study. Breathing rate rose from 18 to 28 and 31 breaths per minute from normal to moderate to severe, an order that held in every test. Recovery took about 3, 10, and 50 seconds, paced by how long the airway-receptor signal lingers in the brainstem, and the circuit followed breathing closely (correlation 0.96–0.98). In WESAD, a spoken stress task raised anxiety and breath depth (*p* < 0.001); speech set the breathing rhythm during the task, and breathing then returned to rest, as the model predicts for healthy people. A compact, provable model links clinical stress measures to the breathing signals that wearables record. It offers measurable targets for monitoring anxiety and depression: how strongly threat drives respiration and how quickly homeostatic regulation restores it.

**Author summary:** Anxiety and depression change the way people breathe: breathing becomes faster, and it takes longer to settle after a stressful moment. Wearables now record breathing continuously, but without a mechanistic account of how the brain’s response to threat shapes each breath, these recordings are hard to interpret. We built a mathematical model that runs from a threat-processing circuit in the amygdala, tuned by a person’s coping capacity and perceived stress, through the brainstem centers that set breathing, to the stretch and irritant sensors in the lungs that feed back on them. Analysis and simulation showed that as the clinical profile worsens, breathing becomes faster both at rest and under threat and recovers more slowly after stress. This recovery is paced not by the threat itself but by how long the brainstem retains signals from the airway sensors. In healthy volunteers performing a public-speaking stress test, speaking itself set the breathing rhythm and masked any stress-driven change in rate, and breathing rate was back at rest once the task ended. Our results identify two quantities that wearables could track, how strongly threat drives breathing and how quickly it recovers, and define the experiments needed to test them in patients.

## Introduction

### Clinical motivation

Anxiety and depressive disorders are among the leading causes of disability worldwide [1]. They develop gradually, as stress load outgrows coping capacity over months and years [2]. As this imbalance consolidates, it reshapes autonomic regulation: resting heart rate (HR) rises, blood-pressure (BP) dynamics shift, and recovery after a stressor slows [3]. Breathing changes early in this progression [4], and it holds a special position among autonomic outputs because it runs under both voluntary cortical control and automatic brainstem control [5]. Emotional states alter the rate and depth of breathing in characteristic ways [6, 7], and anticipatory anxiety alone raises respiratory rate [8]. Psychological stress shifts the balance between voluntary and automatic control toward the automatic brainstem circuits [9], which makes respiratory rate (RR, in breaths per minute) and tidal volume (*V*_*T*_, the air volume of one breath, in liters) sensitive markers of distress. Over-breathing, raised RR, irregular *V*_*T*_, and breathlessness are core features of panic and anxiety that often precede cardiovascular symptoms [10, 11], and spontaneous panic has been proposed to arise from a false suffocation alarm [12]. RR can now be monitored easily in daily life with chest straps or wrist-worn photoplethysmography (PPG) [13], a central element of digital phenotyping in psychiatry [14]. A mechanistic understanding of how anxiety and depression shape respiration would give these recordings direct clinical meaning.

### Research gap

Wearable stress detection, mechanistic physiology modeling, and computational psychiatry each address part of this problem, as reviewed below, and the closest mechanistic account maps clinical questionnaires onto cardiovascular output only [15]. To our knowledge, the present work reports the first model that carries clinically parameterized threat processing all the way to the receptor feedback that sets respiratory rate and depth. Its starting point is the amygdala, whose hyperactivity is a consistent finding across anxiety disorders [16] and which has a functional pathway to the brainstem respiratory network in humans: electrical stimulation of the amygdala inhibits breathing [17]. The coupling runs in both directions, since neurons of the brainstem breathing center project to arousal centers [18]. Such a model has to represent two receptor families. Slowly adapting stretch receptors (SARs) in the airway wall sense lung inflation and brake inspiration through the Hering–Breuer inflation reflex [19, 20], and rapidly adapting receptors (RARs) sense fast airway deformation and accelerate breathing [21, 22]. Both project to the nucleus tractus solitarius (NTS), the brainstem relay that also receives baroreceptor input and passes respiratory signals on to limbic structures including the amygdala [9, 23]. The RAR pathway forms a contains a positive feedback loop: rapid breathing disturbs the airway recruiting further RAR input, and anxiety-driven over-breathing feeds the same loop [10, 22]. Over-breathing also lowers arterial carbon dioxide (CO_2_), which strengthens SAR discharge and the Hering–Breuer reflex [19, 20], and rapid breathing shortens the time available to fill the lungs on each breath [24]. Together these effects set a ceiling on *V*_*T*_ . Recovery matters as much as the peak response, since stress responses in anxiety and depression last longer and return to baseline slowly [2, 25], and recovery carries information that reactivity alone does not [26]. An electrical equivalent circuit offers a compact summary of recovery in the form of one physically interpretable time constant.

### Contributions

Three barriers kept these elements apart until recently. First, respiratory models were built for ventilation and gas exchange: their inputs are chemical, and their parameters are physiological constants tuned for intensive-care and exercise physiology, far from the questionnaires used in mental health. Second, the RAR pathway responds to the rate of change of breathing, so representing its positive feedback requires a model that resolves the breathing pattern dynamically, and steady-state ventilation models average this dynamic away. Third, a threat-processing formulation whose parameters map onto clinical scales, and whose output can drive an autonomic readout, has become available only recently [15]. Three developments now make a mechanistic respiratory model timely: wearables deliver continuous RR in daily life [13]; public datasets such as the Wearable Stress and Affect Detection (WESAD) dataset pair chest-belt respiration with a standardized psychological stressor [27]; and medicine is moving toward patient-specific digital twins, which require compact, mechanistic, and identifiable models [28].

The study examines how clinically parameterized threat processing is translated into respiratory rate, depth, and recovery, and which elements of the respiratory feedback circuitry determine how long breathing stays elevated after stress. Our contributions are as follows.

1. A threat-to-breathing mechanistic model in which clinically parameterized threat input, brainstem drive, SAR braking, RAR acceleration, and a rate-dependent *V*_*T*_ penalty jointly set RR and *V*_*T*_ .
2. A mathematical analysis that proves bounded dynamics and a unique, locally exponentially stable resting state with explicit recovery rates. Together with a slow-manifold reduction showing that, in the moderate and severe regimes, recovery is paced by the decay of RAR memory in the NTS rather than by the threat signal, and an exact peak bound and retention law for the recovery circuit.
3. A physical recovery index: a diode-gated resistor–capacitor (RC) circuit whose discharge time constant condenses recovery into a single value. The null models show that the diode is the element that aligns the circuit’s dynamics with breathing.
4. Multi-level substantiation through numerical verification across 30 noise realizations and four stimulus paradigms, local and global sensitivity analyses. A comparison with the state of the art, and an evaluation on WESAD against five hypotheses specified from the model. All outcomes, including the confounding effect of speech on breathing during stress, are reported in full, and every result is reproducible with our open-source code.

### Related approaches

Three research communities address parts of the problem set out above: wearable stress detection, mechanistic physiology modeling, and computational psychiatry. Table 1 compares the proposed framework with five benchmark approaches from these communities on six attributes: mechanistic structure, threat-input compatibility, respiratory-output generation, recovery-index derivation, alignment with clinical scales, and compatibility with wearable sensing. Each existing approach covers isolated links in the chain from clinical scales to wearable respiratory signals; the proposed model is designed to cover the whole chain.

**Table 1.** Comparison with state-of-the-art approaches.

| Approach | Examples | Mechanistic | Threat input | Respiratory output | Recovery index | Clinical-scale parameters | Wearable-matched output |
| --- | --- | --- | --- | --- | --- | --- | --- |
| Wearable machine-learning stress detection | [27, 29] | – | labels | as feature | – | – | ✓ |
| Integrated cardiopulmonary models | [30, 31] | ✓ | – | ✓ (chemoreflex) | ◦ | – | ◦ |
| Brain–heart interaction models | [32] | ✓ | ◦ | as input | – | – | ✓ (HR) |
| Dynamical-systems mood models | [33] | abstract | ✓ (mood) | – | ✓ | self-report | – |
| Amygdala–cardiovascular model | [15] | ✓ | ✓ | – | ◦ | ✓ | ✓ (HR, BP) |
| This work | – | ✓ | ✓ | ✓ (RR, $V_T$) | ✓ ($\tau$) | ✓ | ✓ (RR) |

✓: included; ◦: partly included; –: outside the scope of the approach. HR: heart rate; BP: blood pressure; RR: respiratory rate; $V_T$ : tidal volume; $\tau$ : discharge time constant of the RC recovery circuit (see Models and methods).

### Wearable machine-learning stress detection

The first community builds stress detectors from wearable data. Machine-learning models trained on heart-rate variability (HRV), skin conductance, and related signals classify stress and mood with high accuracy [29], with binary stress detection on WESAD reaching up to 93% [27]. These detectors treat the physiology between the stressor and the recorded signal as a learned mapping from signals to labels, with the mechanism left implicit. The proposed model complements such detectors by explaining a change in RR as a change in named parameters, such as the threat gain or the recovery rate.

### Mechanistic cardiopulmonary and brain–heart models

The second community builds mechanistic physiology models. Integrated cardiopulmonary models reproduce lung mechanics, gas exchange, and circulation in detail [30], with respiratory control driven by chemoreflexes that respond to arterial CO_2_ and blood acidity (pH) [31]. Classic chemoreflex models explain periodic breathing as an instability of this feedback loop [34], and models of the brainstem respiratory network resolve rhythm generation and its regulation by pulmonary afferents [35]. The proposed model adds the affective route, threat input with receptor feedback, using two respiratory state variables and 14 parameters, a size that keeps parameter estimation from wearable data within reach, as the inference of a participant-specific threat gain from WESAD recordings illustrates. Brain–heart interaction models link central activity to the control of HR, with respiration as a modulating input [32]. The proposed model instead makes respiration an output of central threat processing.

### Dynamical-systems models of mood

The third community, computational psychiatry, relates mental illness to formal models of neural and cognitive computation [36] and, increasingly, of brain–body interactions [37, 38]. Dynamical-systems accounts within it describe mood as a dynamical system with alternative stable states and tipping points, and explain slow recovery with abstract variables [33]. The mathematical analysis presented in (Propositions 3–5) instead gives recovery explicit, measurable time constants.

### Amygdala–cardiovascular model

Conductance-based amygdala models have recently begun to bridge these communities by mapping two clinical questionnaires, the Connor–Davidson Resilience Scale (CD-RISC) for coping [39] and the ten-item Perceived Stress Scale (PSS-10) [40], onto neuronal excitability and cardiovascular output, namely HR and BP [15]. Breathing, the receptor feedback that shapes it, and the dynamics of recovery lie outside their scope. The present model addresses these three elements, and because both models use the same threat signal, they can be combined into one cardiorespiratory model driven by one set of clinical parameters.

## Models and methods

### Model architecture

The model comprises three sequential stages: threat processing, respiratory control, and recovery (Fig 1). The threat stage uses the conductance-based amygdala formulation of [15]; the respiratory control loop, the recovery circuit, and the analysis of both are introduced here. Time *t* is in seconds, except in the threat stage, which runs on a millisecond time scale.

**Fig 1.**
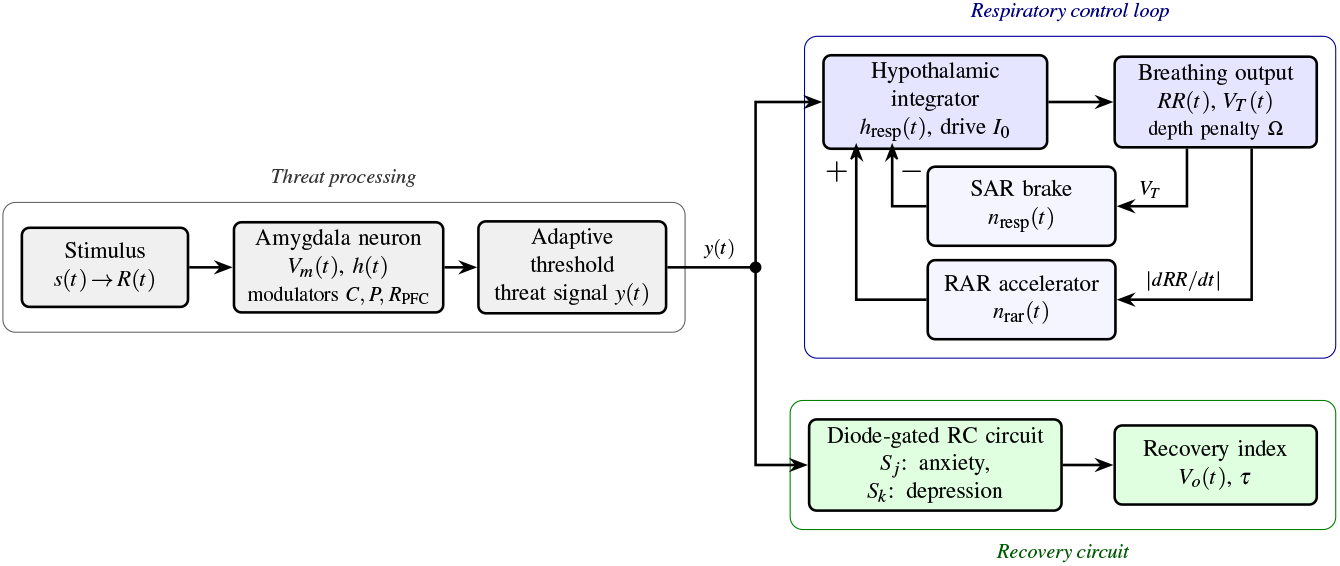
Model architecture. Gray: threat-processing stage, which turns a stimulus *s*(*t*) into the threat signal *y*(*t*) through a conductance-based amygdala neuron with clinically parameterized modulators. The stimulus produces a receptor potential *R*(*t*); the neuron has membrane potential *V*_*m*_(*t*) and slow activation state *h*(*t*); and coping capacity *C*, perceived stress *P*, and prefrontal regulation *R*_PFC_ set its excitability and its adaptive threshold. Blue: respiratory loop, in which the SAR brake lowers (−) and the RAR accelerator raises (+) the hypothalamic drive that sets breathing rate and depth. The hypothalamic respiratory state *h*_resp_(*t*), driven by *y*(*t*) and the constant brainstem drive *I*_0_, sets the respiratory rate *RR*(*t*) and tidal volume *V*_*T*_ (*t*); the depth penalty Ω reduces *V*_*T*_ while RR changes quickly; the SAR brake *n*_resp_(*t*) is driven by *V*_*T*_ ; and the RAR accelerator *n*_rar_(*t*) is driven by |*dRR/dt*|, the magnitude of the rate of change of RR. Green: RC recovery circuit driven by the same threat signal. Switches *S*_*j*_ and *S*_*k*_ connect the anxiety and depression storage capacitors, and the output voltage *V*_*o*_(*t*) decays with the recovery time constant *τ* . The Models and methods section defines every element.

#### Threat input

Consider a stimulus *s*(*t*) ∈ [0, 1] producing a receptor potential *R*(*t*) (mV) that drives a single-compartment, conductance-based amygdala neuron in the Hodgkin–Huxley framework [41]. Its membrane potential *V*_*m*_ (mV) and a slow internal state *h* ∈ [0, 1], which records recent activation, evolve on a millisecond time scale as [15]

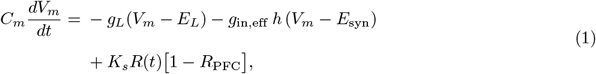

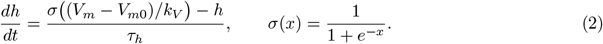

Here *C*_*m*_ is the membrane capacitance; *g*_*L*_ and *E*_*L*_ are the leak conductance and reversal potential. *g*_in,eff_ and *E*_syn_ are the effective inhibitory conductance and synaptic reversal potential. *K*_*s*_ scales the sensory current. *V*_*m*0_ and *k*_*V*_ are the half-activation voltage and slope of the slow state, whose time constant is *τ*_*h*_; and *σ* is the logistic function. Three clinical modulators set the excitability of the neuron: coping capacity *C* ∈ [0, 1] from the CD-RISC, perceived stress *P* ∈ [0, 1] from the PSS-10, and prefrontal regulation *R*_PFC_ ∈ [0, 1], the top-down control exerted by the prefrontal cortex, which gates the sensory drive [42]. Coping and stress enter through *g*_in,eff_ = *g*_in_(1 + 0.4*C* − 0.5*P*), where *g*_in_ is the peak inhibitory conductance. The same modulators set an adaptive threshold *θ*, and a sigmoid of the distance to that threshold gives the threat acknowledgment signal *A* ∈ (0, 1):

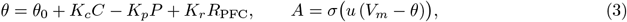

where *θ*_0_ is the baseline threshold, *K*_*c*_, *K*_*p*_, and *K*_*r*_ are threshold coefficients, and *u* is the readout gain. Strong coping and strong prefrontal regulation raise the threshold, and high perceived stress lowers it. Clear threat signals pass to the hypothalamus through the gate

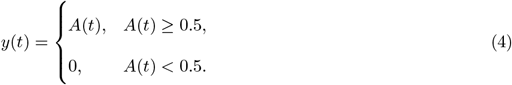

Three settings of the modulators, calibrated in [15], define the normal, moderate, and severe regimes (Table 2). The threat stage is identical in every analysis, so each difference downstream of *y*(*t*) comes from the threat signal or from the respiratory coefficients.

**Table 2.** Model parameters.

| Symbol | Meaning | Normal | Moderate | Severe |
| --- | --- | --- | --- | --- |
| *Threat stage [15]* |  |  |  |  |
| $R_{\text{PFC}}$ | Prefrontal regulation | 0.75 | 0.50 | 0.25 |
| $C$ | Coping capacity | 0.75 | 0.50 | 0.25 |
| $P$ | Perceived stress | 0.25 | 0.50 | 0.75 |
| $\tau_h$ (ms) | Amygdala adaptation | 40 | 100 | 220 |
| *Respiratory loop (rates in $s^{-1}$)* |  |  |  |  |
| $\alpha_{\text{resp}}$ | Threat gain | 0.45 | 0.80 | 1.40 |
| $\alpha_{\text{rar}}$ | RAR gain | 0.20 | 0.25 | 0.30 |
| $\beta_{\text{resp}}$ | SAR brake gain | 0.40 | 0.25 | 0.15 |
| $\gamma_{\text{res}}$ | Passive recovery | 0.30 | 0.15 | 0.08 |
| $\eta_{VT}$ (L) | SAR sigmoid scale | 0.02 | 0.10 | 0.10 |
| $\beta_{\Omega}$ | Depth penalty gain<sup>a</sup> | 0.020 | 0.025 | 0.025 |
| $K_{\text{RAR}}$ | RAR sensitivity<sup>a</sup> | 0.010 | 0.030 | 0.050 |
| $k_n$ | RAR memory gain | 0.50 | 1.00 | 1.50 |
| $\delta_N$ | RAR forgetting rate | 0.40 | 0.10 | 0.02 |

*Shared, threat stage:* $C_m = 1 \mu\text{F}/\text{cm}^2$ ; $g_L = 0.10$ , $g_{\text{in}} = 0.30$ , $K_s = 0.6 \text{ mS}/\text{cm}^2$ ; $E_L = E_{\text{syn}} = -70 \text{ mV}$ ; $V_{m0} = -40 \text{ mV}$ ; $k_V = 8 \text{ mV}$ ; $\theta_0 = -55 \text{ mV}$ ; $K_c, K_p, K_r = 4, 5, 4 \text{ mV}$ ; $u = 0.20 \text{ mV}^{-1}$ . *Shared, respiratory loop:* $RR_{\text{rest}} = g_{rr} = 17 \text{ breaths}/\text{min}$ ; $V_{T,\text{rest}} = g_{VT} = 0.50 \text{ L}$ ; $V_{HB} = 1.0 \text{ L}$ ; $\eta_{hb} = 0.05 \text{ L}$ ; $k_1 = 0.08$ ; $k_2 = 0.48$ ; $q = m = 2$ ; $I_0 = 0.0333 \text{ s}^{-1}$ ; $h_{\text{resp},0} = 0.10$ . <sup>a</sup>Units of $(\text{breaths} \cdot \text{min}^{-1} \cdot \text{s}^{-1})^{-2}$ .

#### Hypothalamic respiratory integrator

The hypothalamic respiratory state *h*_resp_ ∈ [0, 1], a normalized level of respiratory recruitment, integrates descending limbic drive and ascending visceral feedback [9]:

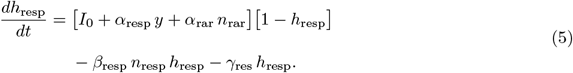

The first bracketed term collects the excitatory drives: the constant brainstem drive *I*_0_ from the medullary rhythm generator [5, 43], the threat input *y* weighted by the threat gain *α*_resp_, and the RAR signal *n*_rar_ weighted by the RAR gain *α*_rar_. The factor (1 − h_resp_) caps recruitment. The negative terms describe inhibition and restoration: the second term is the SAR brake *n*_resp_ with gain *β*_resp_, and the third is passive recovery at rate *γ*_res_. With severity, *α*_resp_ and *α*_rar_ increase while *β*_resp_ and *γ*_res_ decrease (Table 2), describing a system that is highly sensitive to stressors and resistant to baseline return.

#### Stretch-receptor brake

Tidal volume follows the hypothalamic state, but during rapid changes in RR it is transiently reduced, reflecting hypocapnia-induced airway resistance and shortened inspiratory time:

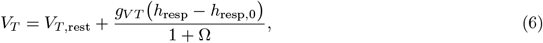

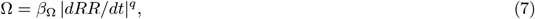

where *V*_*T*,rest_ is the resting tidal volume, *g*_*V T*_ the volume gain, and *h*_resp,0_ the baseline hypothalamic state. The depth penalty Ω has gain *β*_Ω_ and exponent *q*, and *dRR/dt* is evaluated with a backward finite difference. The penalty lumps the two effects of fast over-breathing described in the Introduction, low-CO_2_ enhancement of SAR discharge and shorter inspiratory time, into one rate-dependent term that vanishes at steady state. The SAR output *b*_SAR_ and the NTS brake signal *n*_resp_ are

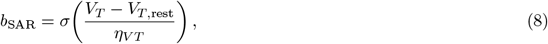

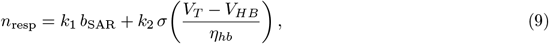

where *η*_*VT*_ and *η*_*hb*_ are the scales of the SAR and Hering–Breuer sigmoids, *V*_*HB*_ is the Hering–Breuer volume threshold, and *k*_1_ and *k*_2_ weight the smooth SAR response and the Hering–Breuer reflex. The reflex engages above about 1–1.7 L in conscious adults [20]. The threshold *V*_*HB*_ = 1.0 L is placed within this range, with *k*_2_*/k*_1_ = 6 so that the reflex dominates once *V*_*T*_ approaches it.

#### Irritant-receptor accelerator

RARs respond to the rate of airway deformation [21, 22]. Because the model operates at the time scale of the breathing pattern, |*dRR/dt*| serves as the proxy for this deformation, and the RAR output *b*_RAR_ and its NTS relay *n*_rar_ are

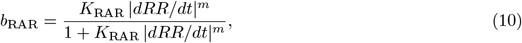

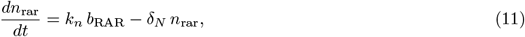

with sensitivity *K*_RAR_, exponent *m*, memory gain *k*_*n*_, and decay rate *δ*_*N*_ . Because *n*_rar_ directly excites *h*_resp_ in Eq (5), faster breathing recruits additional hypothalamic drive, establishing a positive feedback loop unique to the respiratory channel.

#### Breathing output

Respiratory rate is read out from the hypothalamic state as a deviation from rest,

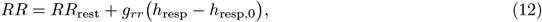

where *RR*_rest_ is the resting rate and *g*_*rr*_ the rate gain. The pair (*RR, V*_*T*_) forms a closed inner feedback loop, since *dRR/dt* enters both Eq (7) and Eq (10) and *V*_*T*_ drives the SAR brake, Eq (9).

#### RC recovery circuit

The circuit topology is shown in Fig 2, the physical meaning of each element in Table 3, and the operating cases in Table 4.

**Table 3.** RC recovery circuit: physical meaning of each element.

| Element | Physical meaning |
| --- | --- |
| $V_{\text{in}} = y$ | Threat signal from the amygdala |
| $R_s$ | Coupling set by coping and perceived stress; low $R_s$ means weak coping and high stress |
| $D$ | One-way loading: threat charges the store, which holds charge between threats |
| $C_j, C_k$ | Stress-load storage linked to anxiety ($S_j$) and depression ($S_k$) |
| $R_L$ | Recovery drain (fixed) |
| $\tau = R_L(S_j C_j + S_k C_k)$ | Discharge time constant (recovery index) |
| $V_o$ | Stored stress load (normalized) |

**Table 4.** RC recovery circuit: operating cases.

| Case | Input regime | $S_j$ | $S_k$ | $\rho = R_s/R_L$ | $\tau$ (s) |
| --- | --- | --- | --- | --- | --- |
| C1 | input off | 0 | 0 | 5 | – |
| C2 | normal | 0 | 0 | 3 | – |
| C3 | moderate | 0 | 1 | 0.2 | 16 |
| C4 | moderate | 1 | 0 | 0.2 | 14 |
| C5 | severe | 1 | 1 | 0.01 | 30 |

**Fig 2.**
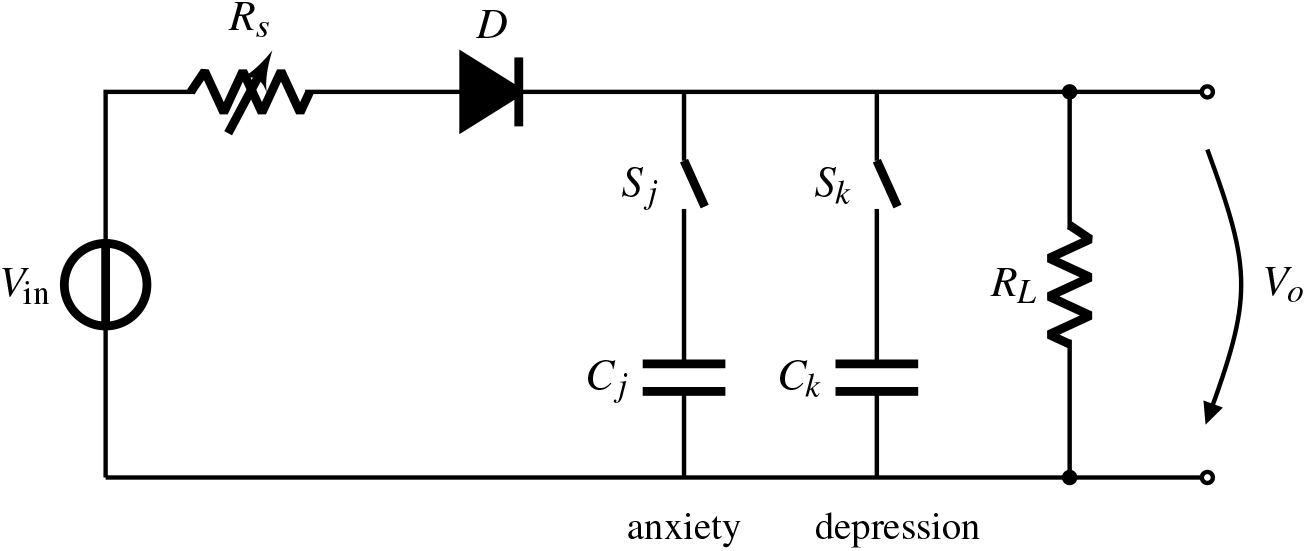
Diode-gated RC recovery circuit. The switches connect the anxiety and depression storage capacitors, the diode admits charge and holds it, and *R*_*L*_ drains it.

The threat signal drives the circuit as its source voltage, *V*_in_ = *y*. Binary switches *S*_*j*_ (anxiety) and *S*_*k*_ (depression), each open (0) or closed (1), connect the storage capacitors *C*_*j*_ and *C*_*k*_. A diode *D* lets current flow in one direction to charge the storage elements and blocks reverse conduction. The stored charge drains through the recovery resistor *R*_*L*_, and the output voltage *V*_*o*_ represents stored stress load. The source resistance *R*_*s*_ encodes the coping-and-stress axis: a high *R*_*s*_ corresponds to strong coping and low perceived stress, and *R*_*s*_ decreases monotonically from the normal to the severe state. With the impedance ratio *ρ* = *R*_*s*_*/R*_*L*_ and the discharge time constant *τ* = *R*_*L*_(*S*_*j*_*C*_*j*_ + *S*_*k*_*C*_*k*_), Kirchhoff’s current law gives, while the diode conducts,

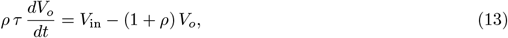

and, while it blocks (*V*_in_ ≤ V_*o*_), *τ dV*_*o*_*/dt* = −V_*o*_. The diode is treated as ideal, and *V*_in_ and *V*_*o*_ are normalized quantities.

### Mathematical analysis

This section establishes properties of the system that hold for all parameter configurations in Table 2; they are verified numerically in the Results. For convenience, let *h* ≡ h_resp_ and *n* ≡ n_rar_ (the amygdala state *h* of Eq (2) does not appear in this section), let (*h*^∗^, n^∗^) denote a steady state, and let *RR*^∗^ be the corresponding resting respiratory rate.

#### Proposition 1

(Bounded dynamics). *For any threat input y*(*t*) ∈ [0, 1], *the set*

*D* = *{*0 ≤ h ≤ 1, 0 ≤ n ≤ *k*_*n*_*/δ*_*N*_ *} is forward invariant. Consequently*

*RR* ∈ [*RR*_rest_ − g_*rr*_*h*_resp,0_, *RR*_rest_ + *g*_*rr*_(1 − h_resp,0_)] = [15.3, 32.3] *breaths/min and V*_*T*_ ≤ V_*T*,rest_ + *g*_*V T*_ (1 − h_resp,0_) = 0.95 *L*.

*Proof*. On *h* = 0, Eq (5) gives *dh/dt* = *I*_0_ + *α*_resp_*y* + *α*_rar_*n* ≥ I_0_ > 0, and on *h* = 1 it gives *dh/dt* = −(β_resp_*n*_resp_ + *γ*_res_) < 0. Since 0 ≤ b_RAR_ < 1, Eq (11) gives *dn/dt* ≥ 0 on *n* = 0 and *dn/dt* = *k*_*n*_(*b*_RAR_ − 1) < 0 on *n* = *k*_*n*_*/δ*_*N*_ . The vector field therefore points into *D* on its boundary, and the bounds on *RR* and *V*_*T*_ follow from Eqs (12) and (6) with Ω ≥ 0.

#### Proposition 2

(Unique resting state). *For a constant input y, the model has exactly one resting state. It satisfies n*^∗^ = 0 *and F* (*h*^∗^) = 0, *where*

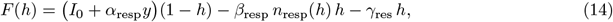

*and n*_resp_(*h*) *is Eq (9) evaluated with* Ω = 0.

*Proof*. At rest *dRR/dt* = 0, so Ω = 0 and *b*_RAR_ = 0, and Eq (11) gives *n*^∗^ = 0; Eq (5) then reduces to *F* (*h*^∗^) = 0. The function *n*_resp_(*h*) increases with *h* through Eqs (6)–(9), so *F* decreases strictly on [0, 1]. Since *F* (0) = *I*_0_ + *α*_resp_*y* > 0 and *F* (1) = −β_resp_*n*_resp_(1) − γ_res_ < 0, *F* has exactly one root.

#### Proposition 3

(Local stability and recovery rates). *With q* = *m* = 2, *the resting state is locally exponentially stable, with eigenvalues* −λ_*h*_ *and* −δ_*N*_, *where λ*_*h*_ = −F ^*′*^(*h*^∗^) > 0 *is the hypothalamic recovery rate. Small deviations therefore decay as RR*(*t*) – 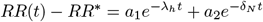, *with constants a*_1_ *and a*_2_ *set by the initial deviation, and the recovery time constant is T*_rec_ = 1*/* min(*λ*_*h*_, *δ*_*N*_).

*Proof*. Since *dRR/dt* = *g*_*rr*_ *dh/dt*, both Ω and *b*_RAR_ are proportional to (*dh/dt*)^2^ near rest and vanish to second order in the deviations (*δh, δn*). The linearization of Eqs (5) and (11) is therefore 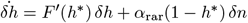 and 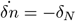 *δn*. This Jacobian is upper triangular with eigenvalues *F* ^*′*^(*h*^∗^) < 0, by Proposition 2, and −δ_*N*_ < 0.

#### Proposition 4

(Slow-manifold recovery). *After the stimulus ends (y* = 0*), let ε* = *δ*_*N*_ */λ*_*h*_(*h, n*) *along the trajectory. If ε* ≪ 1, *then after an initial layer of duration of order* 1*/λ*_*h*_, *h*(*t*) = *H (n*(*t*)) + *O*(*ε*), *where O*(*ε*) *denotes an error of order ε and H*(*n*) *is the unique root of*

*G*(*h, n*) = (*I*_0_ + *α*_rar_*n*)(1 − h) − β_resp_*n*_resp_(*h*)*h* − γ_res_*h. Recovery of RR is then paced by the forgetting rate δ*_*N*_ *of the RAR memory*.

*Proof*. Here *λ*_*h*_(*h, n*) = −∂G/∂h > 0 by the argument of Proposition 2, so *H*(*n*) is a uniformly attracting root of the fast equation. During slow recovery *dRR/dt* is small, so *b*_RAR_ ≈ 0 and *n* decays at rate *δ*_*N*_ . Tikhonov’s theorem on singularly perturbed systems [44] then gives *h* = *H*(*n*) + *O*(*ε*) after the initial layer, so that *RR*(*t*) ≈ RR_rest_ + *g*_*rr*_ *(H*(*n*(*t*)) − h_resp,0_) with 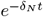.

#### Corollary 1

(Saturation of sensitivity). *At a constant input y* > 0, ∂h^∗^/∂α_resp_ = *y*(1 − h^∗^)/*λ*_*h*_, *which tends to zero as h*^∗^ → 1. *The response to further threat therefore compresses near full recruitment*.

*Proof*. Differentiate *F* (*h*^∗^; α_resp_) = 0 implicitly, using ∂F/∂α_resp_ = *y*(1 − h^∗^) and ∂F/∂h = −*λ*_*h*_.

#### Proposition 5

(Recovery circuit). *Let* 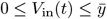, *where the constant* 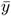 *bounds the input, and V*_*o*_(0) = 0. *Then (i)* 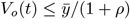 *for all t; (ii) during an interval in which the input equals* 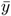, *conduction drives* 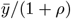 *with charging time constant τ*_*c*_ = *ρτ/*(1 + *ρ*); *(iii) after the input ends at t*_off_,

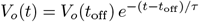 *exactly; and (iv) with both switches open, V*_*o*_ = *V*_in_*/*(1 + *ρ*) *during conduction and the circuit stores zero charge*.

*Proof*. (i) During conduction, Eq (13) gives 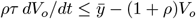, which is negative whenever 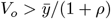, and during blocking *dV*_*o*_*/dt* = −V_*o*_*/τ* ≤ 0. (ii) and (iii) follow by solving the two linear equations, and (iv) follows from Eq (13) with *τ* = 0.

Proposition 5 also explains two circuit properties. Either switch creates storage, so the presence of memory follows OR logic, and parallel capacitances add in *τ*, so its duration is additive. With the diode removed, Eq (13) holds at all times and becomes the first-order low-pass filter *V*_*o*_(*s*)*/V*_in_(*s*) = 1*/*(1 + *ρ* + *ρτs*) in the Laplace variable *s*, which averages an intermittent threat signal over its quiescent intervals. With the diode in place, the network instead acts as a peak-hold circuit that captures and retains transient maxima.

### Simulation protocol

The stimulus consists of five 400-ms pulses of amplitude 0.5 at 0.8, 1.5, 2.2, 2.9, and 3.6 s superimposed on Ornstein–Uhlenbeck noise, a mean-reverting random process with time constant 80 ms and standard deviation 0.12; the sum is clipped to [0, 1], and the stimulus is followed by a 5-s recovery interval [15]. The receptor potential rises toward 40 *s*(*t*) mV with a 15-ms time constant during stimulation and decays with a 50-ms time constant between stimuli. The threat stage is integrated with forward Euler at a 0.1-ms step from *V*_*m*_(0) = −55 mV and *h*(0) = *σ* ((*V*_*m*_(0) − V_*m*0_)*/k*_*V*_), the respiratory loop with forward Euler at a 1-ms step, and the circuit with backward Euler at a 1-ms step. Each regime was simulated over 30 noise realizations (random seeds 42 and 0–28); realization 42, the default seed of the threat-stage reference code, is shown in the figures and is used for the ablations and for the local and global sensitivity analyses. Tables report the mean *±* standard deviation (SD). All regimes start from *h*_resp_ = 0.10, *n*_rar_ = 0, *RR* = 17 breaths/min, and *V*_*T*_ = 0.50 L, nominal adult values. The 10-s window matches the membrane time scale of the threat stage, so the respiratory outputs describe the envelope of the breathing pattern on a compressed time base.

#### Output measures and verification

For each run we record the peak RR, the peak *V*_*T*_, the peak minute ventilation 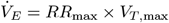 (an upper bound, since the two peaks can occur at different times), the RR at the end of the window, and the resting rate *RR*^∗^ from Eq (14). Regime separation is measured by the peak-RR differences Δ_*NM*_ (normal to moderate), Δ_*MS*_ (moderate to severe), and Δ_*NS*_ (normal to severe). For the circuit we record the peak output and the retention *V*_*o*_(*t*)*/V*_*o*_(*t*_off_) with *t*_off_ = 5 s. The analytical results are checked against simulation: Proposition 2 against a 300-s run with *y* = 0, Proposition 4 by comparing the simulated RR during recovery with *RR*_rest_ + *g*_*rr*_(*H*(*n*_rar_(*t*)) − h_resp,0_), and Proposition 5 by the ratio of the simulated circuit peak to its bound. Numerical accuracy is checked by halving the 1-ms step.

#### Robustness and sensitivity

Robustness is assessed over the 30 realizations and over four stimulus paradigms with 10 realizations each (seeds 0–9): the pulse train, a sustained step of amplitude 0.5 from 0.8 to 4.0 s, a ramp from 0 to 0.7 over the same interval, and noise alone. Two ablations remove the RAR branch (*α*_rar_ = 0) and the depth penalty (*β*_Ω_ = 0). Local sensitivity uses the elasticity *E*_*p*_ = [ΔRR_max_(1.2*p*) − ΔRR_max_(0.8*p*)]*/*[0.4 ΔRR_max_(*p*)] of the peak excursion ΔRR_max_ with respect to each parameter *p*, so that *E*_*p*_ = 0.3 means that a 1% change in *p* changes the excursion by about 0.3%. Because one-at-a-time designs miss interactions [45], a global analysis draws 400 Latin-hypercube samples [46], a stratified design that covers each parameter range evenly, with all 14 respiratory parameters scaled jointly by factors in [0.8, 1.2] [47]. A linear regression of the standardized peak *h*_resp_ on the standardized factors yields standardized regression coefficients *c*_*j*_ and the coefficient of determination *R*^2^.

#### Null models for the circuit

The circuit and the respiratory loop share the input *y*, so part of their agreement comes from the shared input. We therefore compare the Pearson correlation *r* and Spearman rank correlation *r*_*s*_ between *V*_*o*_(*t*) and *RR*(*t*) with those of two null models driven by the same *y*: the same RC network with the diode removed, and a first-order low-pass filter with a 1-s time constant. The coefficients are computed per realization and serve as descriptive measures of shape similarity.

### WESAD analysis

#### Data and preprocessing

WESAD contains chest-belt respiration recorded with a RespiBAN device at 700 Hz from *N* = 15 healthy adults during a 20-min baseline. An amusement condition with funny video clips, the Trier Social Stress Test (TSST) [48], in which participants deliver a speech and perform mental arithmetic aloud before a panel, and guided meditation, together with the six-item short form of the State-Trait Anxiety Inventory (STAI) [49] after each condition [27, 50]. We used the respiration channel as released in compressed form by a WESAD co-author, which subsamples the original signal to 140 Hz, quantizes it to 1% of each channel’s range, and includes the protocol timing [51]. The questionnaire files, which contain the STAI responses and the protocol schedule, were taken from a public copy of the original release [52]; their protocol timings match those of the co-author’s release exactly. The signal was resampled to 25 Hz and band-pass filtered between 0.1 and 0.7 Hz (6–42 breaths/min), and breaths were detected as peaks at least 1.4 s apart with a prominence of 0.35 standard deviations. RR is 60 divided by the breath interval, and breath amplitude, the peak minus the preceding trough, serves as a *V*_*T*_ proxy, normalized to each participant’s baseline median. Breaths within 30 s of a condition boundary were excluded, and each participant was summarized by condition medians. The TSST-related change ΔRR is the TSST median minus the baseline median. As an independent check, a spectral estimate took the dominant frequency of the filtered signal in 60-s windows advanced in 30-s steps within each condition.

#### Hypotheses and statistics

Five hypotheses were specified from the model before the WESAD data were analyzed. H1: the TSST raises RR. H2: breath amplitude rises together with RR. H3: participants with a larger RR rise recover more slowly, measured by fitting 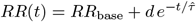, where *RR*_base_ is the participant’s baseline median rate, *d* the amplitude, and 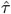 the recovery time constant, from the end of the TSST (*t* = 0) to the next labeled condition (60–300 s), accepting fits with *R*^2^ ≥ 0.3 and *d* > 0. H4 (exploratory): the model-inferred threat gain 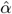, the *α*_resp_ that reproduces a participant’s RR rise at a steady unit threat input with the other moderate-regime parameters, tracks the TSST-related rise in STAI. H5 (exploratory): resting RR is higher in participants with higher baseline STAI, as Proposition 2 predicts across severity. The STAI items (at ease, nervous, jittery, relaxed, worried, pleasant; each 1–4) were scored 20–80 with the three positive items reverse-scored. Paired effects were tested with the Wilcoxon signed-rank test, with effect size *d*_*z*_ (mean paired difference divided by its SD), and associations with *r*_*s*_; *p* denotes the *p*-value of each test. Two analyses added after the pre-specified tests address speech, which drives breathing during the TSST [53, 54]. RR in the silent questionnaire period after the TSST was compared with the identical period after amusement (first 120 s), and RR during the TSST was examined in six successive intervals (0–60, 60–120, 120–180, 180–300, 300–450, and 450–600 s) with Bonferroni correction. Sensitivity to the detector was assessed with prominences of 0.25 and 0.50 and with a 0.08–0.8 Hz band.

### Ethics statement

This study is a secondary analysis of the publicly available, de-identified WESAD dataset [27, 50], which was collected by its original investigators.The dataset is released for academic, non-commercial research only, and the present use, academic research with no commercial purpose, satisfies these terms. The data were analyzed exactly as released: the authors had no contact with participants, had no access to identifying information, and did not attempt to re-identify anyone. Because the study used only anonymized, publicly available data, no separate ethical approval was required, and in the authors’ assessment the analysis posed no risk to participants’ privacy

### Software

All simulation, verification, figure, and WESAD-analysis code (Python with NumPy, SciPy, and Matplotlib) reproduces every number and figure in this paper and is provided as S1 File, under the MIT License, together with a README that maps each script to the corresponding equations, tables, and figures, and the derived numerical outputs. All random draws use fixed seeds, and the simulation results were reproduced exactly with Python 3.12, NumPy 2.4, SciPy 1.17, and Matplotlib 3.10. The WESAD inputs are not redistributed; the code retrieves them from their public sources [51, 52] and also accepts the original release [50].

## Results

Table 5 summarizes the evidence for each main claim; the subsections below give the details.

**Table 5.** Substantiation of the main claims.

| Claim | Evidence | Outcome |
| --- | --- | --- |
| Bounded dynamics | Proposition 1; 30 realizations | $h_{\text{resp}} \in [0.098, 0.938]$, $n_{\text{rar}} \leq 1.60$ |
| One resting state per regime | Proposition 2; 300-s run at $y = 0$ | $RR^*$ agrees within 0.01 breaths/min |
| Recovery paced by RAR memory | Propositions 3 and 4; slow-manifold check | Severe: mean error 0.27 breaths/min |
| Circuit retention and peak | Proposition 5; simulation | $e^{-t/\tau}$ within 0.1 point; C5 at 99% of bound |
| Severity ordering | 30 realizations; 4 paradigms; 400 Latin-hypercube samples | Held in 100% of runs in each test |
| Numerical accuracy | Step halving | Largest change 0.006% |
| Diode aligns $V_o$ with RR | Two null models | $r = 0.96\text{--}0.98$ vs. $0.06\text{--}0.72$ |
| Dominant parameters | Local and global sensitivity | $\alpha_{\text{resp}}$ and $\gamma_{\text{res}}$ |
| Human psychogenic stress | WESAD, $N = 15$; H1–H5; estimator check | STAI and amplitude rise ($p < 0.001$); speech sets TSST rhythm; post-stress RR at baseline |

### Breathing response across severity

Severity ordered both the rate and the depth of breathing (Fig 3, Table 6).

**Table 6.** Simulated respiratory outputs (mean *±* SD, 30 realizations).

| Regime | Peak RR (breaths/min) | Peak $V_T$ (L) | Peak $\dot{V}_E$ (L/min) | RR at 10 s (breaths/min) | Resting $RR^*$ (breaths/min) |
| --- | --- | --- | --- | --- | --- |
| Normal | $18.26 \pm 0.39$ | $0.537 \pm 0.011$ | 9.8 | 17.09 | 16.92 |
| Moderate | $27.91 \pm 0.11$ | $0.818 \pm 0.004$ | 22.8 | 24.31 | 18.20 |
| Severe | $31.14 \pm 0.03$ | $0.916 \pm 0.001$ | 28.5 | 29.50 | 19.95 |

**Fig 3.**
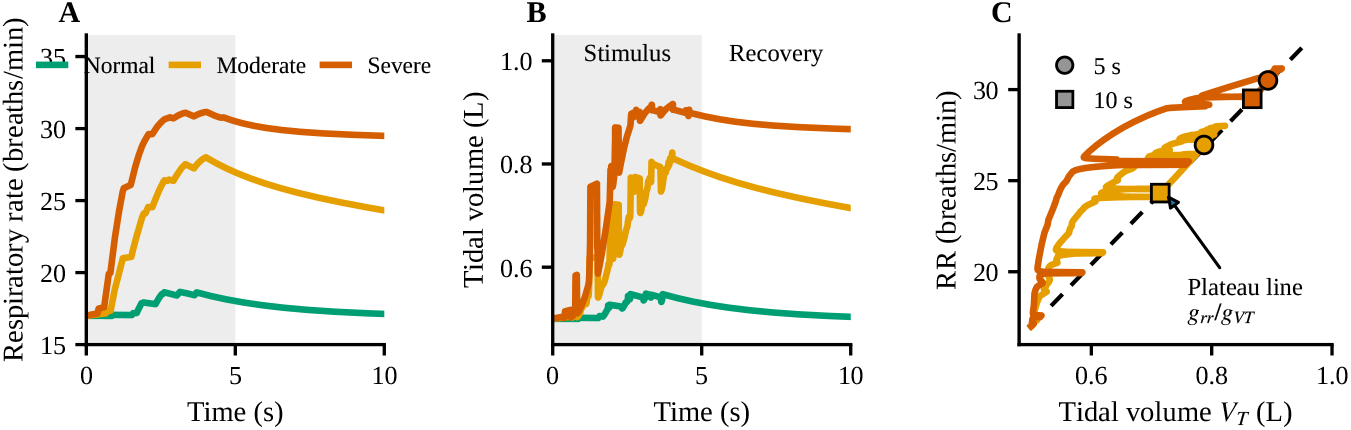
Breathing response across severity (realization 42). A: Respiratory rate. B: Tidal volume. Gray: stimulus interval. C: *V*_*T*_ against RR for the moderate and severe regimes; depth trails rate during acceleration and returns to the plateau line (dashed) once rate is steady. Circles: end of stimulus; squares: end of window.

Peak RR rose from 18.3 breaths/min in the normal regime to 27.9 and 31.1 in the moderate and severe regimes, peak *V*_*T*_ rose from 0.54 to 0.82 and 0.92 L, and peak minute ventilation rose from 9.8 to 22.8 and 28.5 L/min. The step from normal to moderate (9.6 breaths/min) was three times the step from moderate to severe (3.2 breaths/min), as Corollary 1 predicts for a hypothalamic state that reached 0.74 and 0.93, respectively. *V*_*T*_ followed the same 3:1 pattern because at every peak

ΔRR/Δ*V*_*T*_ = *g*_*rr*_*/g*_*V T*_ = 34 breaths*·*min^−1^·L^−1^. During acceleration, *V*_*T*_ trailed RR as the depth penalty acted, peaking at Ω = 3.3 and 9.6 in the moderate and severe regimes, and returned to the plateau line once RR was steady (Fig 3C). Peak rates of 28–31 breaths/min and minute ventilation of 23–29 L/min correspond to acute over-breathing, as during panic episodes [10]. The hypothalamus works like a throttle with a maximum setting: stronger threat input and weaker braking push it further open, and close to full throttle extra drive adds little, so the severe regime sits closer to the moderate regime than the moderate regime sits to normal. Rate and depth follow the same throttle and rise together, and when breathing speeds up quickly each breath has less time to fill the lungs, so depth trails rate until the rate settles. The model therefore predicts quick, shallow breaths at the start of an acute stress response, followed by deeper breaths as the response stabilizes.

### Feedback pathways, resting state, and recovery

The SAR induced inhibition grew lung inflation in all regimes (Fig 4A). Because peak *V*_*T*_ (0.92 L) stayed below *V*_*HB*_, as Proposition 1 guarantees, the Hering–Breuer term acted through the lower part of its sigmoid (*σ* = 0.16) and contributed as much as the smooth SAR term (*k*_2_*σ* = 0.075 against *k*_1_*b*_SAR_ = 0.079). The RAR output fired in bursts during the acceleration of RR and returned to zero once RR was steady (Fig 4B). The NTS integrated these bursts to 0.72 and 1.57 in the moderate and severe regimes (Fig 4C).

**Fig 4.**
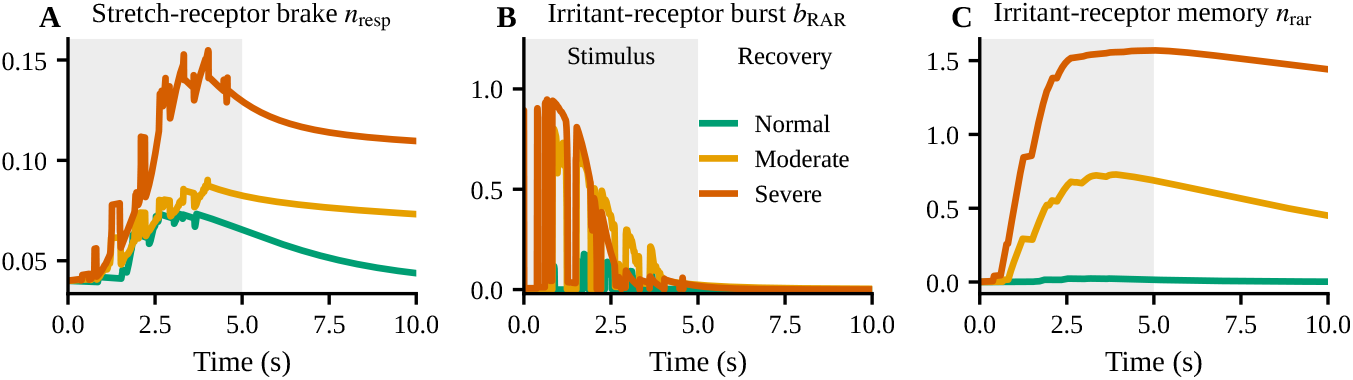
Feedback pathways (realization 42). A: SAR brake *n*_resp_, which grows with lung inflation. B: RAR burst *b*_RAR_, active during the acceleration of breathing. C: NTS memory *n*_rar_ of those bursts, which fades slowly in the severe regime.

Each regime crossed *F* (*h*) = 0 once (Fig 5A), at resting rates of 16.9, 18.2, and 19.9 breaths/min, and a 300-s run at *y* = 0 converged to the same values within 0.01 breaths/min. The hypothalamic recovery rates were *λ*_*h*_ = 0.37, 0.20, and 0.13 s^−1^, and with *δ*_*N*_ = 0.4, 0.1, and 0.02 s^−1^ Proposition 3 gives recovery time constants of 2.7, 10, and 50 s. During recovery in the severe regime, the time-scale ratio was *ε* = 0.03 and the slow-manifold prediction matched the simulated RR with a mean error of 0.27 breaths/min (Fig 5B). In the moderate regime, *ε* = 0.31 and the mean error was 1.4 breaths/min, as expected for weaker time-scale separation. The stretch receptors act as a brake that tightens as the lungs inflate, and the irritant receptors act like a motion sensor that fires as breathing speeds up. The NTS keeps a memory of that firing, and Proposition 4 shows that in the severe regime this memory, which fades over about 50 s, sets the pace of recovery. A brief airway sensation thus keeps breathing elevated long after it ends. The resting set point also rises with severity, so the model predicts faster breathing at rest in more severe states, in the direction of the baseline over-breathing reported in panic disorder [55]. Resting RR and the post-stress decay rate from a wearable therefore carry information about the braking, recovery, and memory parameters of the model.

**Fig 5.**
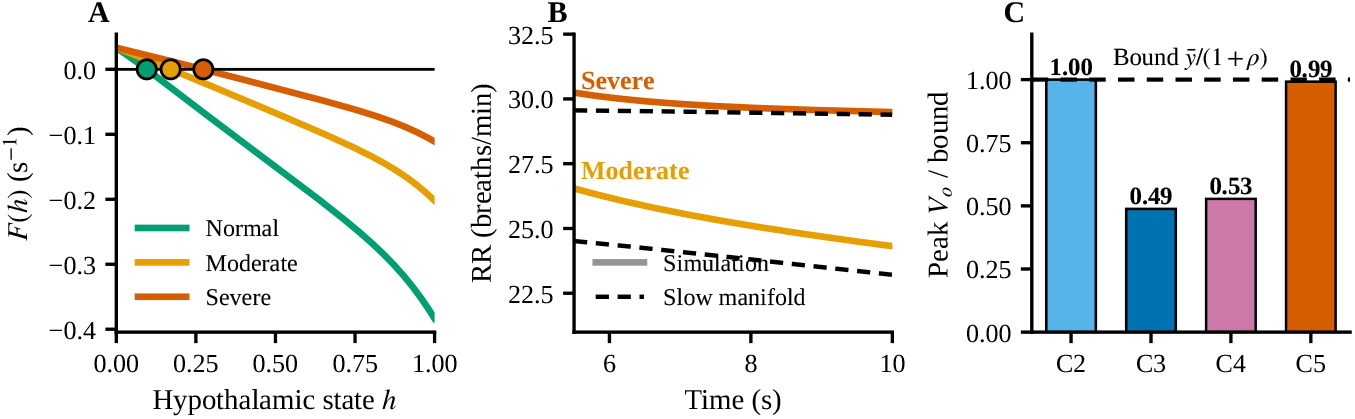
Verification of the mathematical analysis. A: The function *F* (*h*) of Eq (14) at *y* = 0; each regime crosses zero once, at its resting state (circles, Proposition 2). B: RR during recovery (solid) and the slow-manifold prediction of Proposition 4 (dashed). C: Peak circuit output divided by the bound 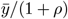 of Proposition 5, maximum over 30 realizations.

### RC recovery circuit

Case C1 stayed at zero because its source is off, and C2 followed the brief normal-regime threat signal through the divider of Proposition 5(iv), reaching 0.18, and returned to zero after each pulse (Fig 6A). Cases C3, C4, and C5 charged with *τ*_*c*_ = 2.7, 2.3, and 0.30 s to peaks of 0.39, 0.42, and 0.98. For C5, the severe-regime threat signal reaches 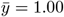, so the bound of Proposition 5 is 0.990, and because *τ*_*c*_ = 0.30 s is shorter than the 0.4-s pulses, the output attained 98.8% of the bound on average (99.2% at most). C3 and C4, with *τ*_*c*_ several times longer than a pulse, reached about half of their bounds (Fig 5C). After the stimulus ended, C3, C4, and C5 retained 73.2%, 70.0%, and 84.7% of their output at 5 s, matching *e*^−t/τ^ to within 0.1 percentage point (Fig 6B). C5 has *τ* = 30 s, the sum of 14 and 16 s.

**Fig 6.**
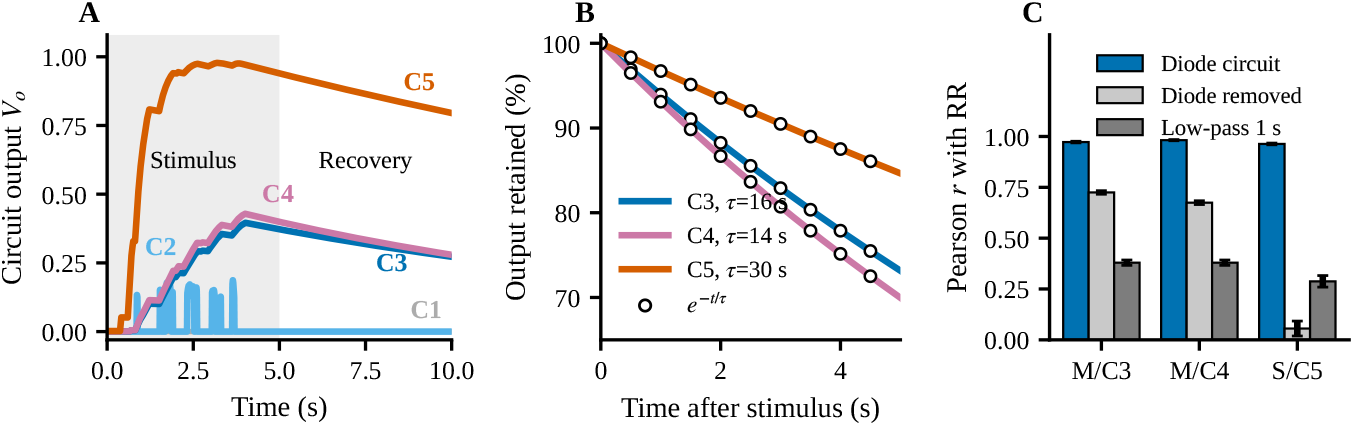
RC recovery circuit (realization 42). A: Output *V*_*o*_(*t*) for cases C1–C5 of Table 4. B: Output retained after the stimulus ends; circles show *e*^−t/τ^ . C: Pearson correlation with RR for the diode circuit and two null models (mean *±* SD over 30 realizations; M: moderate, S: severe).

The circuit output correlated with RR at *r* = 0.973 *±* 0.004 (moderate, C3), 0.982 *±* 0.003 (moderate, C4), and 0.963 *±* 0.004 (severe, C5), with *r*_*s*_ = 0.92, 0.95, and 0.90. With the diode removed, *r* fell to 0.72, 0.67, and 0.06, and a 1-s low-pass filter of *y* reached 0.38, 0.38, and 0.29 (Fig 6C). The capacitor is a reservoir of stress load: weak coping and high stress (low *R*_*s*_) let threat fill it quickly, the diode keeps the load in it between threats, and *R*_*L*_ drains it. Anxiety and depression each add a reservoir, and together they double the storage and the recovery time, in line with the greater severity and chronicity of comorbid anxiety and depression [56]. The null models show why the diode matters: between threat pulses the hypothalamic drive holds breathing high, the diode holds the circuit in the same way, and the diode-free network, a low-pass filter, averages the pauses away. The time constant *τ* is therefore a single recovery index that can be fitted to the decay of a wearable signal after a stressor.

### Robustness and sensitivity

The ordering normal < moderate < severe held in all 30 realizations for peak RR, peak *V*_*T*_, and the circuit peaks (Fig 7A), with smallest gaps of 9.0 and 3.1 breaths/min and coefficients of variation of 2.1%, 0.4%, and 0.1%. It also held in every realization of all four paradigms (Fig 7B): peak ΔRR was 1.3, 11.0, and 14.2 breaths/min for pulses, 1.9, 12.2, and 14.4 for the step, 1.8, 11.5, and 14.4 for the ramp, and 0.0, 2.8, and 12.1 for noise alone. Halving the time step changed no output by more than 0.006%. Removing the RAR branch reduced Δ_*NS*_ by 4.1%, from 12.49 to 11.98 breaths/min, and removing the depth penalty changed it by 0.1%.

**Fig 7.**
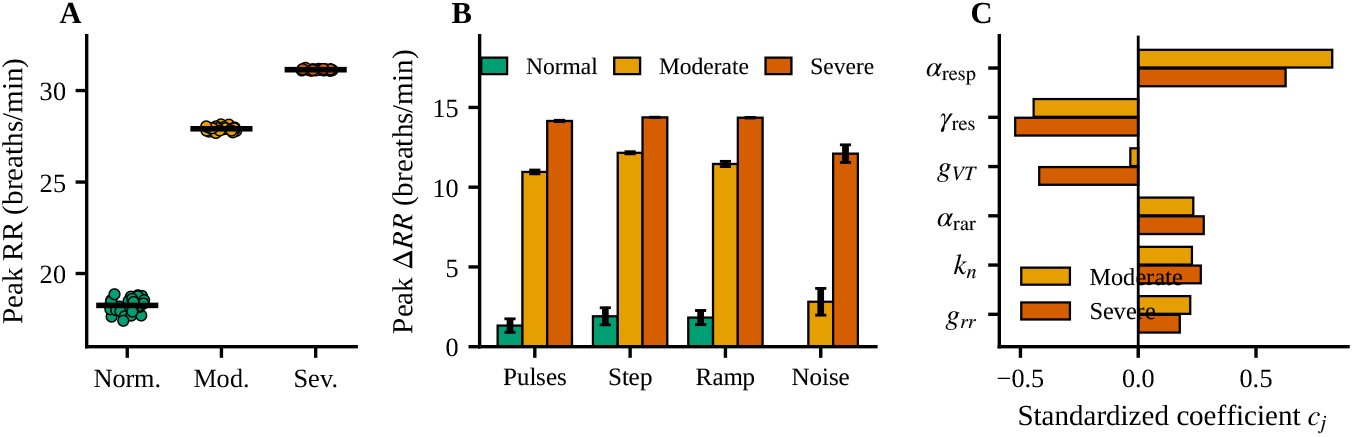
Robustness and sensitivity. A: Peak RR in each of 30 noise realizations; bars mark the mean. B: Peak ΔRR under four stimulus paradigms (mean *±* SD, 10 realizations each). C: Standardized regression coefficients *c*_*j*_ of the six most influential parameters on peak hypothalamic drive (400 joint *±*20% samples).

Locally, excluding the output gain *g*_*rr*_, whose elasticity is close to 1 by construction because the excursion is proportional to it (1.08 moderate, 1.02 severe), *α*_resp_ had the largest elasticity (0.32 moderate, 0.06 severe), followed by *γ*_res_ (−0.17, −0.05). All elasticities were smaller in the severe regime, as Corollary 1 predicts, except those of *g*_*VT*_ and *k*_2_. The depth and Hering–Breuer parameters, which matter more as *V*_*T*_ approaches the reflex threshold. Globally, the ordering held in all 400 samples, with a smallest moderate-to-severe gap of 2.1 breaths/min, and the standardized regression coefficients (*R*^2^ = 0.99 and 0.97) ranked *α*_resp_ first (0.82 moderate, 0.63 severe), then *γ*_res_ (−0.44, −0.52), then *α*_rar_ and *k*_*n*_ (0.23–0.28), with *g*_*VT*_ (−0.42) entering in the severe regime (Fig 7C). The ordering belongs to the structure of the model, since it survives changes in noise, stimulus form, feedback components, and all parameters jointly. The noise-only paradigm is especially telling: background fluctuations alone raised breathing by 12.1 breaths/min in the severe regime, because its lowered threshold lets everyday noise register as threat, a model counterpart of heightened reactivity to ambiguous cues in anxiety. Two levers dominate, the threat gain and the passive recovery rate, and they correspond to two treatment targets: lowering threat drive through prefrontal regulation and cognitive therapy, and speeding recovery through interventions such as slow-breathing training [4].

### Test against human psychogenic stress (WESAD)

The TSST was effective as a psychological stressor: STAI rose from 36.0 *±* 6.7 at baseline to 61.8 *±* 7.0 after the TSST (*p* < 0.001, *n* = 15 participants with complete questionnaires). Table 7 summarizes the pre-specified tests.

**Table 7.**
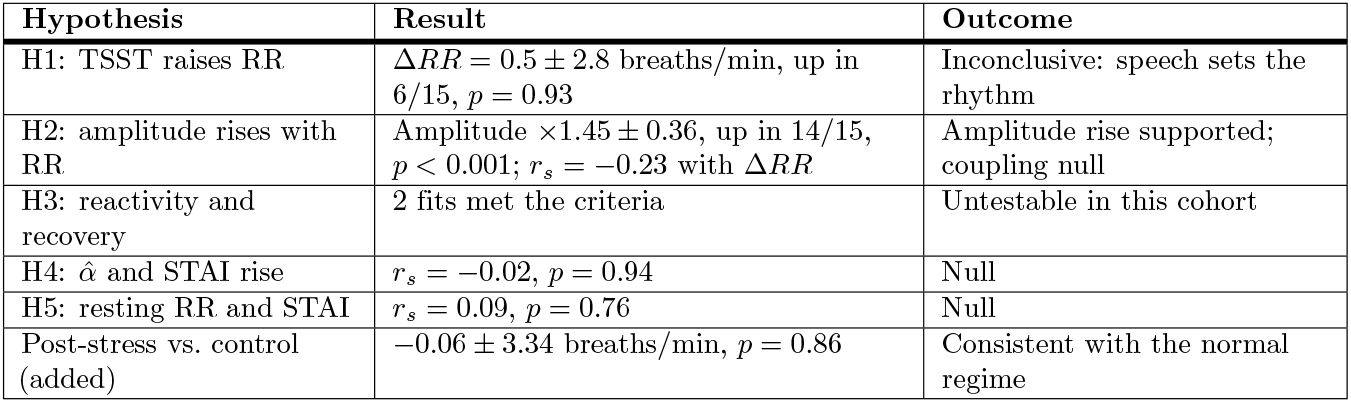
WESAD hypothesis tests (*N* = 15).

| Hypothesis | Result | Outcome |
| --- | --- | --- |
| H1: TSST raises RR | $\Delta RR = 0.5 \pm 2.8$ breaths/min, up in 6/15, $p = 0.93$ | Inconclusive: speech sets the rhythm |
| H2: amplitude rises with RR | Amplitude $\times 1.45 \pm 0.36$, up in 14/15, $p < 0.001$; $r_s = -0.23$ with $\Delta RR$ | Amplitude rise supported; coupling null |
| H3: reactivity and recovery | 2 fits met the criteria | Untestable in this cohort |
| H4: $\hat{\alpha}$ and STAI rise | $r_s = -0.02$, $p = 0.94$ | Null |
| H5: resting RR and STAI | $r_s = 0.09$, $p = 0.76$ | Null |
| Post-stress vs. control (added) | $-0.06 \pm 3.34$ breaths/min, $p = 0.86$ | Consistent with the normal regime |

RR was 16.7 *±* 1.9 breaths/min at baseline and 17.2 *±* 2.8 during the TSST, a change of 0.5 *±* 2.8 breaths/min that was positive in 6 of 15 participants (*p* = 0.93, *d*_*z*_ = 0.17). Breath amplitude during the TSST was 1.45 *±* 0.36 times baseline and higher in 14 of 15 participants (*p* < 0.001), and the changes in amplitude and RR were weakly and negatively associated (*r*_*s*_ = −0.23, *p* = 0.41) (Fig 8B and 8C). Amusement raised RR to 18.3 *±* 1.8 breaths/min (*p* = 0.010) and guided meditation lowered it to 10.7 *±* 2.1 (*p* < 0.001). Because RR changed little after the TSST, only 2 participants met the recovery-fit criteria, which leaves H3 untestable in this cohort. For the same reason the inferred threat gain had median (range) 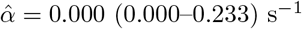, and its association with the STAI rise was null (*r*_*s*_ = −0.02, *p* = 0.94; H4), as was the association of baseline RR with baseline STAI (*r*_*s*_ = 0.09, *p* = 0.76; H5). Every detector setting gave the same picture: the TSST change in RR ranged from −0.4 to +1.0 breaths/min (smallest *p* = 0.25), the amplitude ratio from 1.42 to 1.51, and the amplitude–rate association from *r*_*s*_ = −0.34 to −0.17.

**Fig 8.**
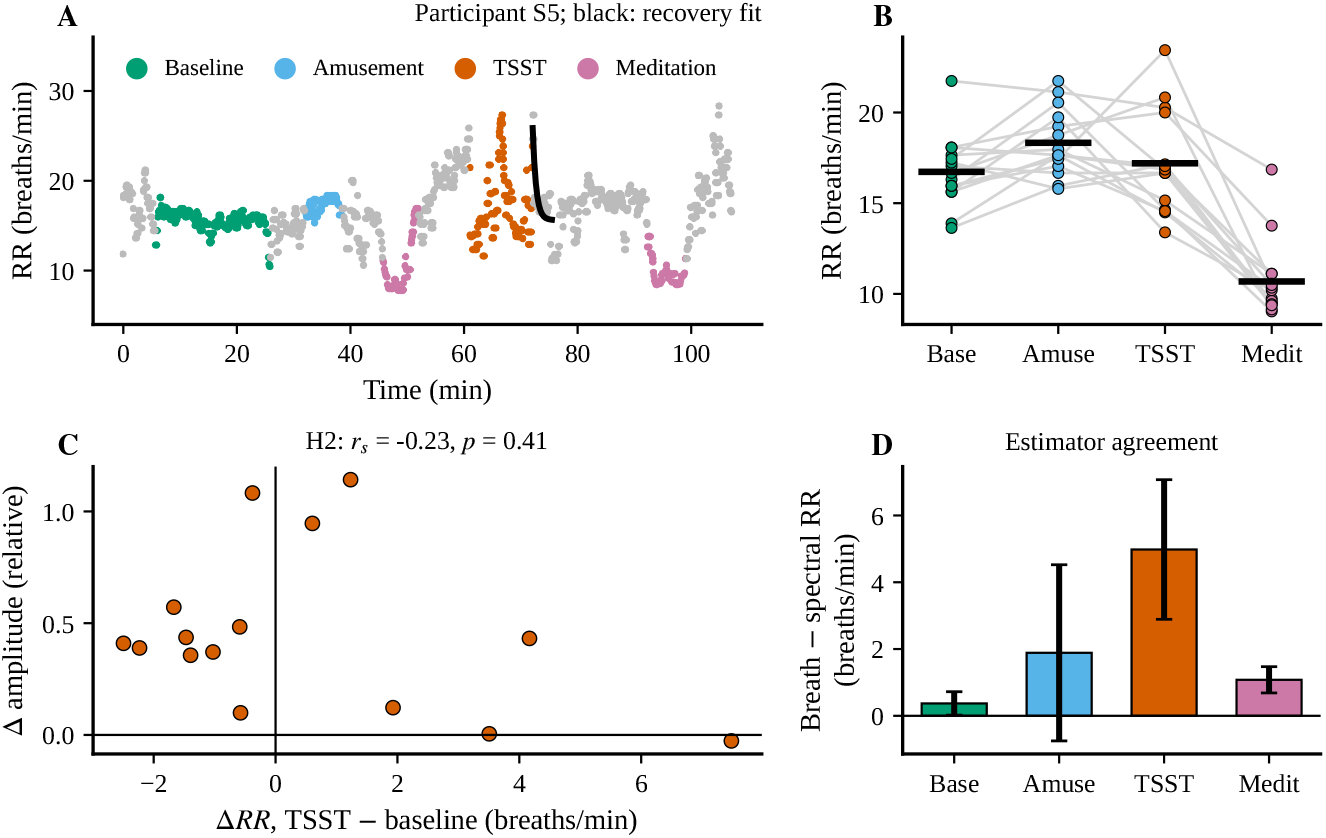
WESAD evaluation. A: Breath-by-breath RR of one participant across the protocol, colored by condition (gray: transitions and reading tasks); black: fitted recovery after the TSST. B: RR per participant by condition; bars mark the means. C: TSST change in RR against the change in breath amplitude (H2). D: Breath-by-breath minus spectral RR by condition (mean *±* SD); the large TSST difference is the signature of speech breathing.

The validity analysis explains the rate result (Fig 8D). At baseline, breath-by-breath and spectral RR agreed within 0.37 *±* 0.35 breaths/min, which confirms the detector, and they agreed within 1.1 *±* 0.4 during meditation and 1.9 *±* 2.6 during amusement, where laughter perturbs breathing. During the TSST, the two estimates differed by 5.0 *±* 2.1 breaths/min, with a spectral RR of 12.2 breaths/min against 16.4 at baseline. This is the signature of speech breathing, in which short inspirations alternate with long phonatory expirations whose timing follows the utterance [53, 54]. The difference stayed at this level from the first minute of the TSST onward, so speech occupies the whole stress condition. Across six successive intervals, one (300–450 s) showed a nominal rise of +1.9 breaths/min (*p* = 0.010), above the Bonferroni threshold of 0.008. In the silent questionnaire period after the TSST, RR was 0.30 *±* 2.65 breaths/min relative to baseline (*p* = 0.64) and matched the identical period after amusement (difference −0.06 *±* 3.34 breaths/min, *p* = 0.86, *n* = 14).

WESAD confirms two elements of the model and marks the boundary of a third. The stressor raised state anxiety strongly, and breath amplitude rose in 14 of 15 participants, the depth component of the model’s respiratory response, with part of that rise reflecting the deep inspirations of speech. The rate component is measurable only outside speech, and in the silent period after the stressor these healthy participants breathed at their baseline rate, consistent with the normal regime of the model: a small peak response (Table 6) and fast recovery (Proposition 3). The moderate and severe predictions, larger rate rises with slow, memory-paced recovery, therefore call for clinical cohorts and speech-free stressors, as the Discussion outlines, and the estimator agreement used here offers a simple quality gate that separates autonomic from speech-driven breathing in such studies.

## Discussion

Three biological insights emerge from the analysis. First, recovery after a stressor is governed by a memory rather than by the stressor itself: in the moderate and severe regimes, breathing returns to rest at the forgetting rate *δ*_*N*_ of the airway-receptor signal held in the NTS (Propositions 3 and 4), so a brief airway sensation can keep breathing elevated for tens of seconds after the threat signal has ended. This places a candidate substrate for the prolonged physiological activation described in anxiety and depression [25, 26] in a visceral afferent pathway rather than in limbic threat processing alone. Second, respiratory recruitment saturates (Corollary 1), so the step from normal to moderate is three times the step from moderate to severe. Peak breathing rate therefore separates mild from moderate states better than moderate from severe, whereas resting rate and recovery time continue to change across the whole range (16.9, 18.2, and 19.9 breaths/min; 2.7, 10, and 50 s). Third, the WESAD analysis shows that during a spoken stressor speech controls the respiratory rhythm, so respiratory rate under the most widely used laboratory stressor is not a clean readout of autonomic drive; the divergence between breath-by-breath and spectral estimates of RR identifies this situation.

The results show that one compact model, driven by clinical measures of coping and stress, reproduces an ordered set of breathing responses, predicts a severity-dependent resting state, and condenses recovery into physical time constants, and that each property is proved analytically and confirmed numerically. This gives respiratory wearables a mechanistic vocabulary: a rise in resting RR maps to weaker braking or slower recovery, a slow decay after a stressor maps to a small *δ*_*N*_ or a large *τ*, and a large response to a mild threat maps to a high *α*_resp_. The analysis also identifies the two parameters that dominate the response, which are the natural targets of psychological and breathing-based interventions, and it supplies the respiratory half of a cardiorespiratory digital twin whose parameters can be estimated from everyday recordings [28].

Four next steps follow from the present results. First, the WESAD evaluation shows what an external test of the rate predictions requires: a stressor that leaves breathing free, such as silent mental arithmetic or driving, a cohort spanning the severity range, and minute-scale recordings to which *δ*_*N*_ and *γ*_res_ can be fitted through Proposition 4, since the 10-s protocol, set by the membrane time scale of the threat stage, compresses time. Second, the model and the circuit each have one resting state (Propositions 2 and 5) and represent persistence through slow relaxation; adding a regenerative feedback branch would create two stable states and allow direct tests of tipping-point behavior [33]. Third, PPG provides HR, HRV, and RR from one wrist sensor [13], which enables joint estimation of cardiovascular and respiratory parameters. Fourth, interventions that calm the stress response, such as slow-breathing training [4, 57], enter the model as a rise in *R*_*s*_ or *γ*_res_, so the model can serve as a testbed for closed-loop therapy.

## Conclusion

This work introduced, to our knowledge, the first model that carries clinically parameterized threat processing to respiratory rate and tidal volume through explicit stretch-receptor and irritant-receptor feedback. A mathematical analysis proved bounded dynamics, a unique and locally stable resting state that rises with severity, and a slow-manifold law under which recovery is paced by the forgetting rate of airway-receptor memory, with time constants of 2.7, 10, and 50 s across regimes. A diode-gated RC circuit condensed recovery into one time constant, obeyed an exact peak bound and retention law, and tracked breathing because its diode holds charge between threats. The severity ordering held across noise, stimulus form, ablation, and joint parameter variation. In WESAD, a spoken stressor raised state anxiety and breath amplitude, speech set the breathing rhythm during the stressor, and healthy adults returned to their baseline rate afterward, consistent with the normal regime; clinical cohorts with speech-free stressors are the next test of the moderate and severe predictions. Together, these results link clinical scales, brain circuits, and wearable breathing signals in one analyzable and reproducible framework for patient-specific digital twins in anxiety and depression.

## Supporting information

**S1 File. Code, derived results, and documentation**. Python code that reproduces all simulations, the verification of Propositions 1–5, the robustness and sensitivity analyses, Figs 3–8, and the WESAD analysis, together with the derived numerical outputs, a README, a requirements file listing package versions, and the MIT License (ZIP).

## Acknowledgments

The authors are thankful to the Walton Institute for Information and Communication Systems Science, South East Technological University, Ireland.

## Author contributions

**Conceptualization:** Malik Faizan, P. J. White, Indrakshi Dey

**Data curation:** Malik Faizan, Indrakshi Dey

**Formal analysis:** Malik Faizan, P. J. White, Indrakshi Dey

**Investigation:** Malik Faizan, P. J. White, Indrakshi Dey

**Methodology:** Malik Faizan, P. J. White, Indrakshi Dey

**Software:** Malik Faizan, Indrakshi Dey

**Supervision:** P. J. White, Indrakshi Dey

**Visualization:** Malik Faizan, Indrakshi Dey

**Writing – original draft:** Malik Faizan, P. J. White, Indrakshi Dey

**Writing – review & editing:** Malik Faizan, P. J. White, Indrakshi Dey

